# High ionic strength vector formulations enhance gene transfer to airway epithelia

**DOI:** 10.1101/2024.01.22.576687

**Authors:** Ashley L. Cooney, Laura Marquez Loza, Kenan Najdawi, Christian M. Brommel, Paul B. McCray, Patrick L. Sinn

**Affiliations:** University of Iowa, Department of Pediatrics; Iowa City, IA 52242, USA; University of Iowa, Center for Cystic Fibrosis Gene Therapy; Iowa City, IA 52242, USA; University of Iowa, Department of Microbiology and Immunology, Iowa City, IA 52242, USA

## Abstract

A fundamental challenge for cystic fibrosis (CF) gene therapy is ensuring sufficient ransduction of airway epithelia to achieve therapeutic correction. Hypertonic saline (HTS) is frequently administered to people with CF to enhance mucus clearance. HTS transiently disrupts epithelial cell tight unctions, but its ability to improve gene transfer has not been investigated. Here we asked if increasing the concentration of NaCl enhances the transduction efficiency of three gene therapy vectors: adenovirus, AAV, and lentiviral vectors. Vectors formulated with 3-7% NaCl exhibited markedly increased transduction for all hree platforms, leading to anion channel correction in primary cultures of human CF epithelial cells and enhanced gene transfer in mouse and pig airways *in vivo*. The mechanism of transduction enhancement nvolved tonicity but not osmolarity or pH. Formulating vectors with a high ionic strength solution is a simple strategy to greatly enhance efficacy and immediately improve preclinical or clinical applications.

**One Sentence Summary:** Formulating adenoviral, AAV, and lentiviral vectors with hypertonic saline remarkably enhances lung gene transfer. (114 characters, including spaces)

## INTRODUCTION

Cystic fibrosis (CF) is an autosomal recessive disease caused by pathogenic variants of the cystic fibrosis ransmembrane conductance regulator (*CFTR*) gene. CFTR protein conducts Cl^-^ and HCO^3-^ anions at the apical surface of epithelial cells. Decreased activity or loss of CFTR in the airways leads to a series of complications including bacterial colonization, chronic inflammation, and mucus accumulation. Numerous studies have established that *CFTR* complementation restores anion transport both *in vitro* and *in vivo*^1–3^. The advent of the small molecule therapy of elexacaftor, tezacaftor, and ivacaftor (ETI), trade name Trikafta, demonstrated that restoring CFTR improves lung function and quality of life in people with CF. Although ETI treatment benefits the majority of people with CF, there remains a pressing need to develop treatments for all people with CF. Gene therapy by *CFTR* complementation can restore anion channel activity regardless of the mutation. Despite numerous clinical trials (reviewed in^4^), no CF gene therapies have advanced past Phase II trials due, in part, to low efficacy of CFTR gene transfer.

The conducting airways are challenging to transduce using topically applied gene therapy reagents due to an enormous surface area and multiple host defense mechanisms to resist viral vector uptake^4^. Vehicle formulations such as lysophosphatidylcholine (LPC)^2, 5^, EGTA^6, 7^, methylcellulose^8^, and perflurocarbon^9^ mprove viral vector transduction and subsequent transgene expression in cultured cells and animal models; however, none of these examples are currently FDA approved for pulmonary administration. The ideal vector formulation that enhances gene transfer would be FDA approved and cost effective. Hypertonic saline (HTS) is commonly aerosolized at 3-7% as a topical lung treatment for people with CF to improve mucus clearance. HTS transiently disrupts tight junctions^10^ and exhibits multiple actions including: a mucolytic agent (disrupting mucus gel), an expectorant (hydrolyzing airway surface liquid), a mucokinetic agent (promoting cough-mediated clearance), and increasing ion flux of two thiols (glutathione and thiocyanate) that protect against oxidative injury (reviewed in^11^). Thus, HTS hydrates surface epithelia, breaks ionic bonds, dislodges mucus, and enhances mucociliary clearance^12^. Clinical benefits and safety track records of his treatment are well established for people with CF.

Here we screened reagents approved for topical airway delivery. We were interested in reagents which decreased transepithelial resistance to increase access to basolateral receptors and enhance gene transfer. We formulated adenoviral (Ad) vectors, adeno-associated viral (AAV) vectors, or the lentiviral vector human mmunideficiency virus (HIV) with 1-7% NaCl and applied the mixtures to the apical surface of primary human airway epithelial cells (HAE). We found that increasing the NaCl tonicity of the vector formulation correlated with enhanced gene transfer in a dose dependent fashion and functionally restored CFTR anion ransport *in vitro*. Focused entry studies with Ad suggest that vector transduction improves with increasing onic strength, remains receptor-mediated, and requires a low pH endosomal step. Finally, we asked if vectors delivered in hypertonic saline enhanced gene transfer *ex vivo* and *in vivo*. Ad and AAV transduction was significantly increased in pig explants *ex vivo*. When we delivered Ad with NaCl to both mouse (7% NaCl) and pig airways (5% NaCl), we observed significantly enhanced gene transfer *in vivo*. Additionally, we aerosolized GP64 pseudotyped HIV lentivirus formulated in 5% NaCl to pig airways and observed widespread reporter gene expression greater than the isotonic saline control. Together, these findings indicate hat formulating viral vectors with hypertonic saline for gene addition or gene editing approaches may be a simple means to improve transduction.

## RESULTS

### Not all vehicles that decrease transepithelial resistance enhance gene transfer

Disrupting the epithelial cell tight junctions allows Ad5-based viral vectors access to their basolaterally localized coxsackie and adenovirus receptor (CXADR)^13^. Unless otherwise specified, these studies use the Ad5 serotype. We screened several compounds for their ability to reduce the transepithelial electrical resistance (TEER) of HAE and improve transduction of an Ad vector. First, reagents were applied to the apical surface of HAE for 5 minutes and then TEER was measured using an ohmmeter. Consistent with previous reports, lysophosphatidylcholine (LPC) (0.1%) and EGTA (250 mM) reduced TEER and increased gene transfer^2, 6^. The clinically approved reagents doxorubicin (5 μM), NaCl (3.6%), and surfactant (Infasurf, 35 mg/ml) all decreased TEER (**Supplemental Fig. 1A**). In separate cultures, Ad-GFP (MOI=250) in 50 µl of DMEM combined 1:1 with the test reagents was applied to the apical surface for 2 hours. 5 days post-reatment, cells were imaged by fluorescence microscopy. Doxorubicin, surfactant, and 0.9% NaCl (PBS) did not enhance Ad gene transfer. Of note, the 3.6% NaCl formulation (mixed 1:1 from 7.2%) conferred remarkable transduction (**Supplemental Fig. 1B**). Based on this initial observation, this concentration of NaCl was selected for the many of the subsequently described Ad-based studies.

### Increasing saline tonicity enhances gene transfer in a dose and time dependent manner

To determine if increasing saline tonicity further improves Ad transduction in HAE, we performed a dose-response of 1-7% NaCl formulation (final concentration, **Table 1**) with Ad-GFP (MOI=250). We observed a dose-dependent increase in transduction efficiency with increasing NaCl tonicity (**Fig. 1A, B**). Basal, secretory, and ciliated cells were all transduced in a dose-dependent manner (**Fig. 1C, Supplemental Fig. 2A**). HAE permeability to dextran also increased in the presence of 4-7% NaCl (**Fig. 1D**), which correlated with increasing GFP expression (**Fig. 1E**). The highest concentrations of NaCl (5-7%) reduced cell viability as determined by flow cytometry (**Supplemental Fig. 2B**) and increased LDH release (**Supplemental Fig. 2C**). To visualize that cell surface integrity and ciliation remained intact, we performed scanning electron microscopy (SEM) on HAE that were untreated or treated with NaCl (3.6%) for 2 hours and observed no overt physical differences compared to the no NaCl control (**Supplemental Fig. 2D**). The therapeutic range of NaCl delivered to patients by aerosolization is up to 7%^14^; however, based on the cell viabilty results, we restricted the delivered concentration of NaCl to 3.5-4.5% for subsequent *in vitro* studies.

**Fig. 1.**
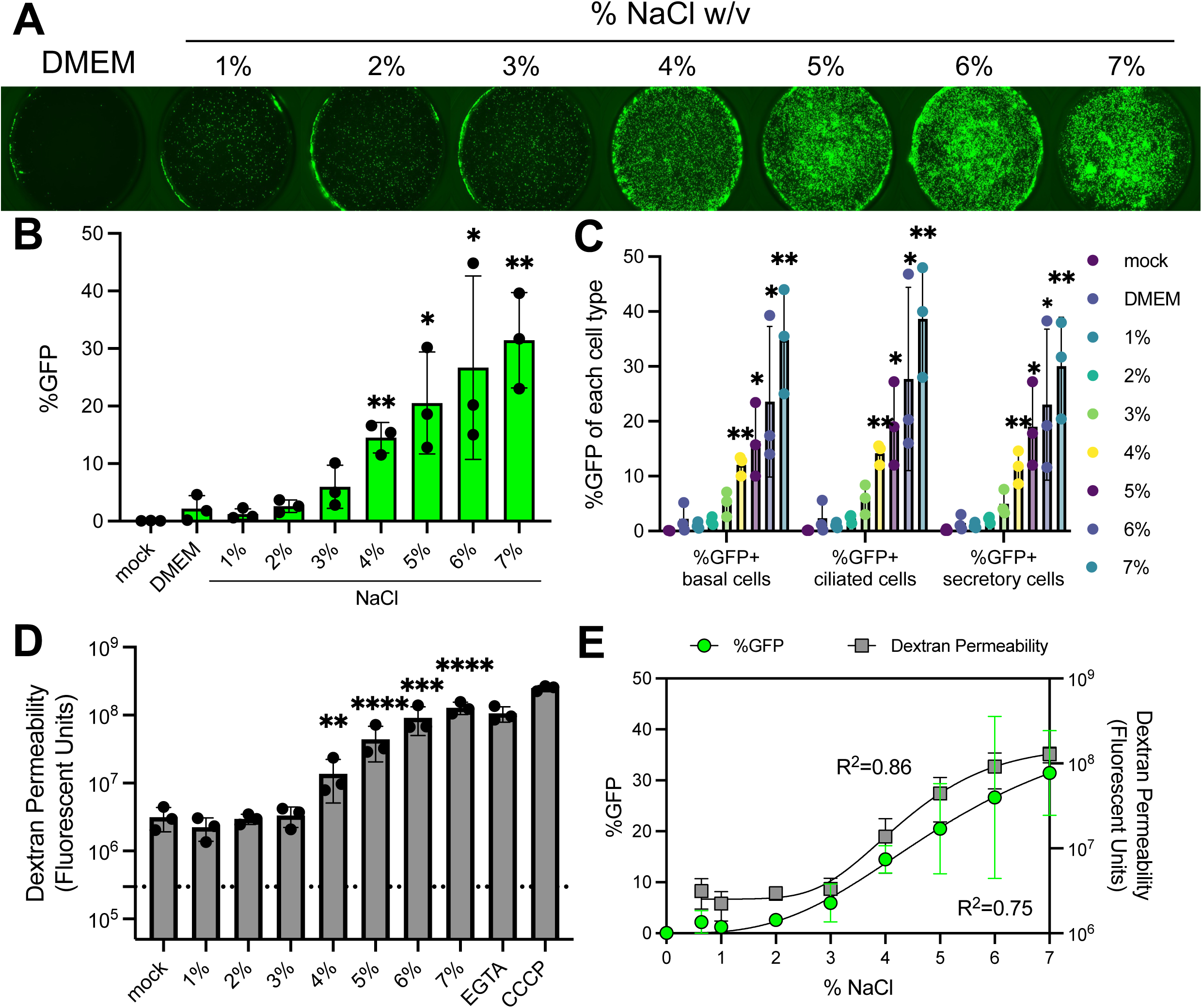
Increasing saline tonicity enhances apical Ad gene transfer in airway epithelia. (**A**) Ad-CMV-eGFP (MOI=250) was co-delivered to primary human airway epithelial cultures with DMEM or NaCl ranging from 1-7% (final concentration) and imaged 5 days post-transduction. 0.33 cm^2^ transwells were imaged at 2X. (**B**) Percentage of GFP+ cells in each condition quantified by flow cytometry. (**C**) Percentages of GFP+ cells by cell type from 1-7% NaCl. (**D**) Tight junction permeability was measured by the passage of dextran from the apical to basolateral media. Airway epithelial cells were left untreated or treated with NaCl (1-7%) for 2 hours. Following the treatment, apical dextran was applied and basolateral media was collected 30 minutes later and measured for fluorescent units were measured using a plate reader. (**E**) Correlative %GFP (R^2^=0.75). from Fig. 1B and Dextran Permeability (R^2^=0.86) from Fig. 1D were plotted together. N=3, *p<0.05, **p<0.005, ***p<0.0005, ****p<0.00005. Statistical differences were determined by one-way ANOVA.

**Table 1:** Molarity and osmolarity of NaCl, KCl, mannitol, and NMDG up to 7%. Tonicity, molarity, and osmolarity of DMEM and 1-7% NaCl, KCl, mannitol, and N-methyl-D-glucamine gluconate (NMDG). Hypertonic indicated a NaCl solution >0.9% DMEM has additional inorganic salts (i.e., CaCl_2_, MgSO_4_, KCl, and NaHCO_3_) and is considered isotonic.

| *Tonicity* | *Isotonic* | *Hypertonic* |  |  |  |  |  |  |
| --- | --- | --- | --- | --- | --- | --- | --- | --- |
|  | **DMEM** |  |  |  |  |  |  |  |
| **% NaCl** | 0.64% | 1% | 2% | 3% | 4% | 5% | 6% | 7% |
| **Molarity** | 110 mM | 170 mM | 340 mM | 510 mM | 680 mM | 860 mM | 1.03 mM | 1.2 M |
| **Osmolarity** | 219<br>mOsmol/L | 342<br>mOsmol/L | 684<br>mOsmol/L | 1,027<br>mOsmol/L | 1,369<br>mOsmol/L | 1,711<br>mOsmol/L | 2,053<br>mOsmol/L | 2,693<br>mOsmol/L |
| **% KCl** | 0.4% | 1% | 2% | 3% | 4% | 5% | 6% | 7% |
| **Molarity** | 50 mM | 130 mM | 270 mM | 400 mM | 540 mM | 670 mM | 800 mM | 940 mM |
| **Osmolarity** | 107<br>mOsmol/L | 268<br>mOsmol/L | 537<br>mOsmol/L | 805<br>mOsmol/L | 1,073<br>mOsmol/L | 1,341<br>mOsmol/L | 1,610<br>mOsmol/L | 1,878<br>mOsmol/L |
| **%**<br>**mannitol** |  | 1% | 2% | 3% | 4% | 5% | 6% | 7% |
| **Molarity** |  | 55 mM | 110 mM | 170 mM | 220 mM | 270 mM | 330 mM | 380 mM |
| **Osmolarity** |  | 55<br>mOsmol/L | 110<br>mOsmol/L | 170<br>mOsmol/L | 220<br>mOsmol/L | 270<br>mOsmol/L | 330<br>mOsmol/L | 380<br>mOsmol/L |
| **% NMDG**<br>**gluconate** |  | 1% | 2% | 3% | 4% | 5% | 6% | 7% |

**Table 1: Molarity and osmolarity of NaCl, KCl, mannitol, and NMDG up to 7%.**
| **Molarity** |  | 50 mM | 100 mM | 150 mM | 200 mM | 260 mM | 310 mM | 360 mM |
| --- | --- | --- | --- | --- | --- | --- | --- | --- |
| **Osmolarity** |  | 342<br>mOsmol/L | 684<br>mOsmol/L | 1,027<br>mOsmol/L | 1,369<br>mOsmol/L | 1,711<br>mOsmol/L | 2,053<br>mOsmol/L | 3,693<br>mOsmol/L |

We next asked how the length of exposure to NaCl or Ad impacted transduction in cultured HAE. As depicted schematically (**Supplemental Fig. 3A**), HAE were incubated with constant Ad-GFP for 120 min and varied times for NaCl (3.6%) at 5, 15, 30, 60, or 120 min. In parallel, HAE were incubated with constant NaCl (3.6%) and varied Ad-GFP incubation times. GFP expression was visualized 5 days post-transduction and quantified by flow cytometry (**Supplemental Fig. 3B**). The %GFP+ cells increased incrementally with ncreasing temporal exposure to either Ad-GFP or NaCl, with significant increase at 1 hour Ad-GFP with constant NaCl (3.6%) exposure and maximal expression after a 2 hour co-treatment. Given these time-dependent results, we established a standard 2 hour co-delivery protocol.

### NaCl-mediated Ad transduction restores anion transport

We hypothesized that the NaCl conferred increase in transduction efficiency would enhance Ad-CFTR complementation of anion transport in CF airway epithelia. We next co-delivered Ad-CFTR formulated with NaCl (3.6%) to the apical surface of primary CF HAE for 2 hours. Five days post-delivery, bioelectric properties were measured in Ussing chambers as previously reported^15^. Without NaCl (3.6%) formulation, no change in Cl^-^ conductance was observed in response to the CFTR agonists forskolin and IBMX (F&I) or CFTR inhibitor GlyH in cultured primary CF epithelia treated with Ad-CFTR (**Fig. 2C**). In contrast, cells that received Ad-CFTR and NaCl (3.6%) showed wild-type levels of Cl^-^ secretion as measured by short circuit current (**Fig. 2A, B**) and conductance (**Fig. 2C, D**). Parallel CF airway cultures were transduced as ndicated to determine transgene expression by microscopy (**Fig. 2E**) and measured by flow cytometry (**Fig. 2F**) or western blot (**Fig. 2F, inset**). These results suggest that NaCl formulation of Ad-CFTR improved gene ransfer and conferred wild-type levels of CFTR mediated anion transport in CF epithelial cells.

**Fig. 2.**
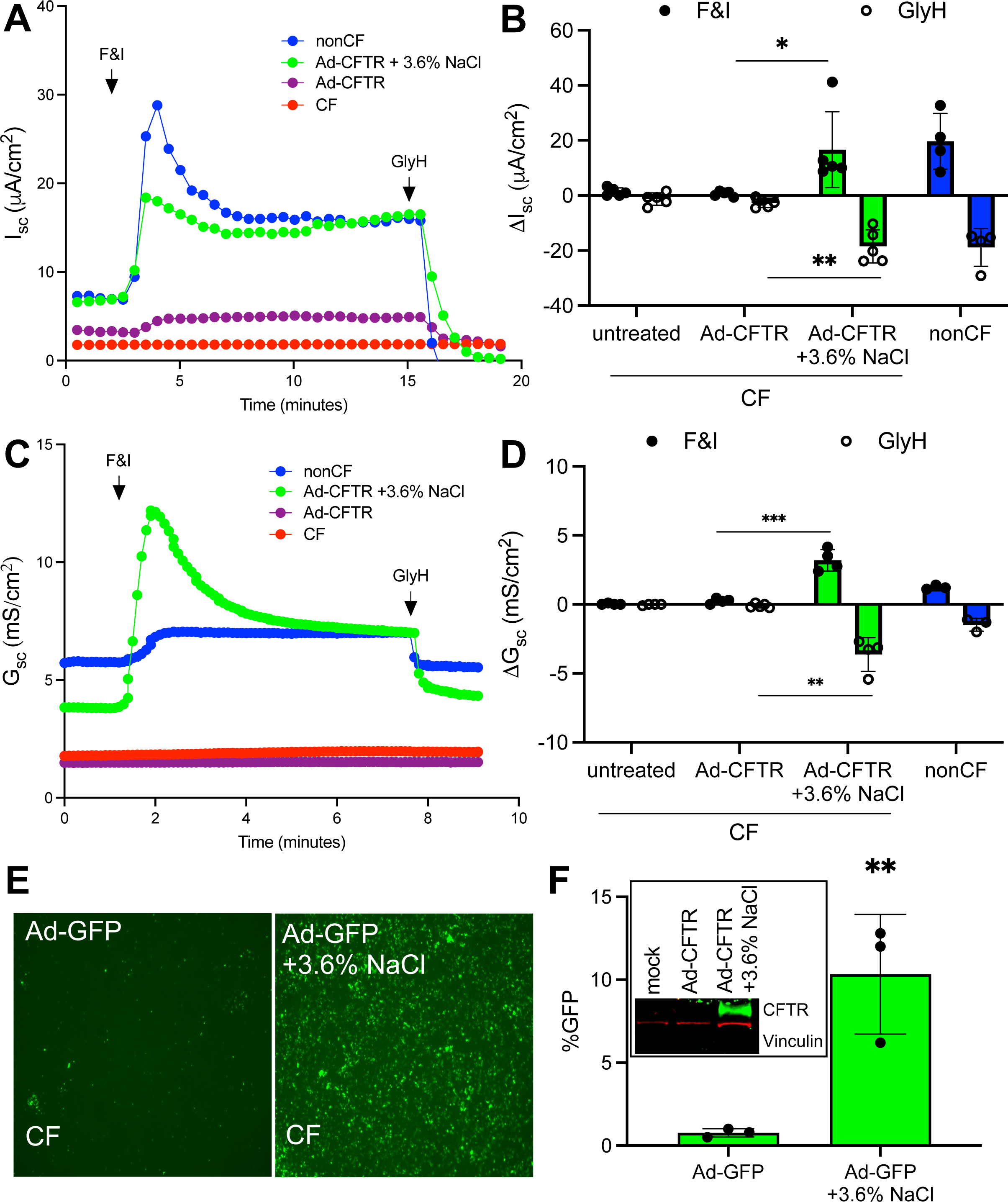
Apically applied Ad-CFTR + 3.6% NaCl restores anion transport defect in CF airway epithelia. (**A, B**) Short circuit current measurements of CF airway epithelia alone or treated with Ad-CFTR or Ad-CFTR + 3.6% NaCl (MOI=250) compared to non-CF. Responses to Forskolin and IBMX (F&I) and GlyH-101 (GlyH) are reported. (**C, D**) Short circuit conductance in response to F&I and GlyH N=5. (**E**) Representative images of parallel CF cultures transduced with Ad-GFP (MOI=250) and (**F**) quantified by flow cytometry (N=3). Inset: representative western blot of CF HAE untreated or transduced with Ad-CFTR or Ad-CFTR + 3.6% NaCl. Western blot was probed for CFTR (green) and Vinculin (red) (loading control). *p<0.05, **p<0.005, ***p<0.0005. Statistical differences were determined by Student’s t-test or one-way ANOVA.

### Impact of osmolarity, pH, and ionic strength on NaCl enhanced gene transfer

To investigate possible mechanisms for the enhanced transduction by NaCl formulations, we compared 3.6% NaCl to the ionic osmolytes 3.6% NaC_6_H_11_O_7_ (NaGluconate), 3.6% KCl, 0.9% NaCl, and 0.9% NaCl with he non-ionic osmolyte mannitol of equivalent osmolarity to the NaCl (3.6%) formulation (**Table 2**). The ndicated regents were co-delivered with Ad-GFP apically to HAE for two hours. Microscopy and flow cytometry quantification was performed five days later. NaCl (3.6%) conferred the greatest increase in ransduction efficiency (**Fig. 3A, B**), whereas equivalent tonicity (3.6%) or osmolarity (620 mM, 1,230 mOsmo/L) with ionic or non-ionic osmolytes did not significantly enhance gene transfer. We next determined if the pH of NaCl formulation impacts transduction. Given that the unadjusted pH of 3.6% NaCl n deionized water is ∼6, we adjusted the pH of the NaCl (3.6%) solution to 5, 6, 7, 7.4, and 8. Following Ad-GFP delivery, and flow cytometry quantification protocols in HAE, no significant differences in %GFP+ cells were observed across this pH range (**Fig. 3C**). To investigate another potential mechansim, we asked if onic strength was responsible for enhanced vector transduction. We compared KCl (lower ionic strength han NaCl for a given percent solution), mannitol (uncharged sugar) and N-methyl-D-glucamine gluconate (charged sugars) at 1-7% tonicities (**Table 1** and **Fig. 3D**). We observed dose-dependent increases with KCl and N-methyl-D-glucamine gluconate whereas mannitol did not enhance gene transfer (**Fig. 3E-G**). The formulation was not dependent on osmolarity or pH, but ionic strength increased the Ad transduction efficiency.

**Fig. 3.**
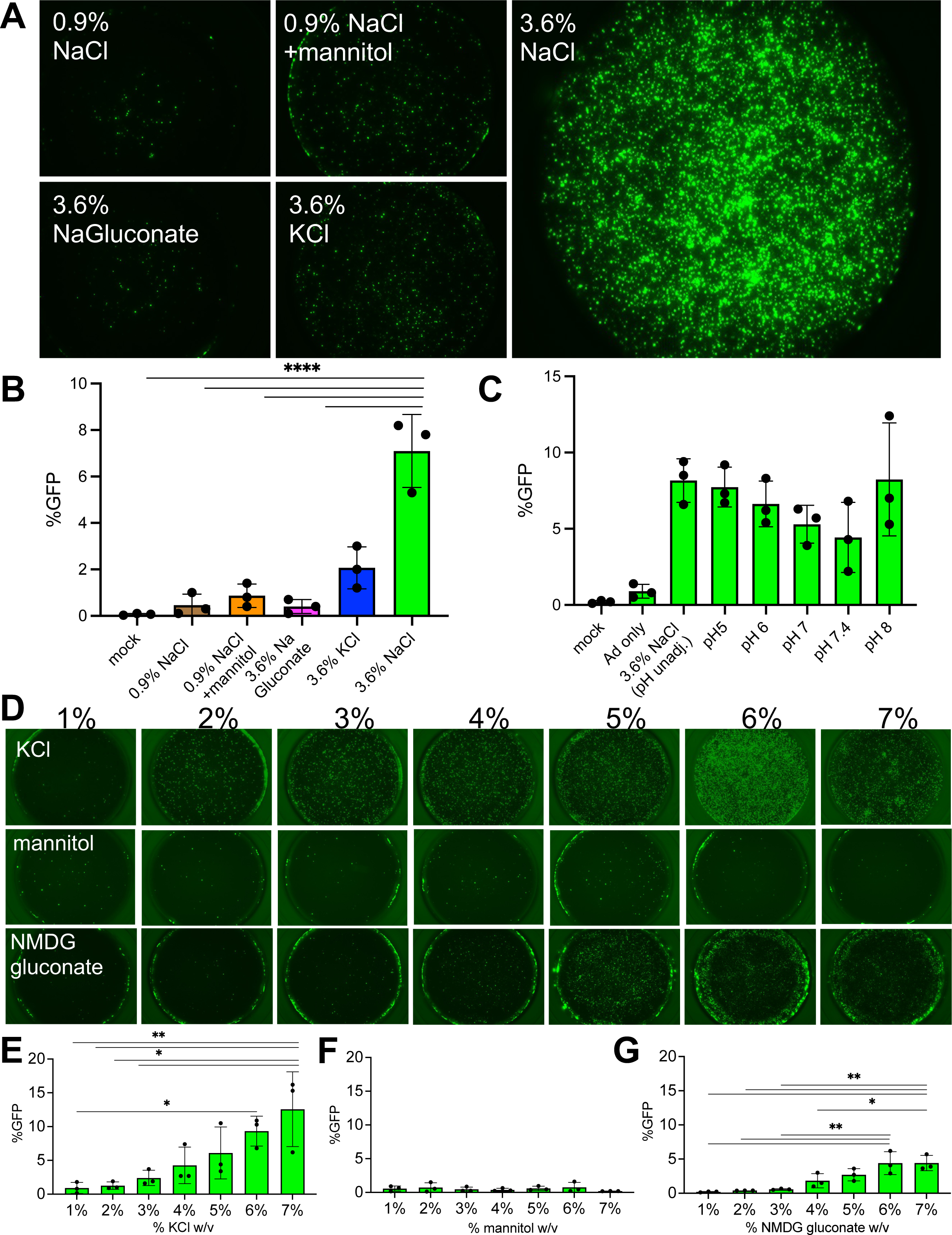
Entry is dependent on ionic strength but not osmolarity, osmolytes, or pH. (**A**) Isotonic NaCl (0.9%), Isotonic NaCl + the non-ionic osmolyte mannitol with equivalent osmolarity (620 mM, 1,230 mOsmol/L) to 3.6% NaCl, NaGluconate (3.6%), and KCl (3.6%) were co-delivered apically to HAE with Ad-GFP (MOI=250) for 2 hours. (**B**) 5 days post-transduction, cells were imaged and GFP expression was quantified by flow cytometry. (**C**) Ad-GFP was co-delivered with NaCl (3.6%) at pH 5, 6, 7, 7.4 and 8 for 2 hours. 5 day post-transduction GFP was quantified by flow cytometry. (**D**) KCl, mannitol, or N-methyl-D-glucamine (NMDG) gluconate 1-7% was formulated with Ad-GFP (MOI=250) and applied to the apical surface of HAE for 2 hours. Cultures were imaged 5 days post-transduction and GFP expression was quantified by flow cytometry for (**E**) KCl, (**F**) mannitol, or (**G**) NMDG gluconate. N=3, *p<0.05, ****p<0.00005. Statistical differences were determined by one-way ANOVA.

**Table 2.** Molarity and osmolarity of ionic and non-ionic osmolytes. Molecular weight, tonicity, molarity and osmolarity for NaCl (0.9% and 3.6%), mannitol, NaGluconate, and KCl are presented.

|  | **3.6% NaCl** | **0.9% NaCl** | **0.9% NaCl**<br>**+mannitol** | **3.6%**<br>**NaGluconate** | **3.6% NaCl** |
| --- | --- | --- | --- | --- | --- |
| **Molecular weight** | 58.44 g/mol | 58.44 g/mol | Mannitol:<br>182.172 g/mol | 590 g/mol | 74.55 g/ml |
| **Tonicity (%)** | 3.6% | 0.9% | 11.2% | 3.6% | 3.6% |
| **Molarity** | 620 mM | 150 mM | 620 mM | 61 mM | 480 mM |
| **Osmolarity** | 1,230<br>mOsmol/L | 310 mOsmol/L | 1,230<br>mOsmol/L | 120 mOsmol/L | 970 mOsmol/L |

### NaCl-mediated gene transfer remains receptor dependent and requires a low pH endosome

Next, we investigated whether the NaCl enhanced Ad5 transduction is receptor dependent, using CRISPR/Cas9 RNPs to knockout the Ad5 receptor CXADR in primary human airway basal cells^16–18^. The electroporated primary cells were seeded on transwell membranes and cultured at an air-liquid interface until well-differentiated (∼21 days). CXADR^KO^ cells and donor-matched airway epithelia were transduced with Ad5-GFP and Ad21-GFP in the presence or absence of NaCl (3.6%). Ad21 enters airway cells through a CXADR-independent mechanism using CD46 as a cellular receptor^19^. Ad5 + NaCl did not transduce CXADR^KO^ epithelia, whereas Ad21 + NaCl readily transduced CXADR^KO^ cells (**Fig. 4A, B**). Following receptor engagement and internalization, Ad5 transduction requires escape from low pH endosomes.

**Fig. 4.**
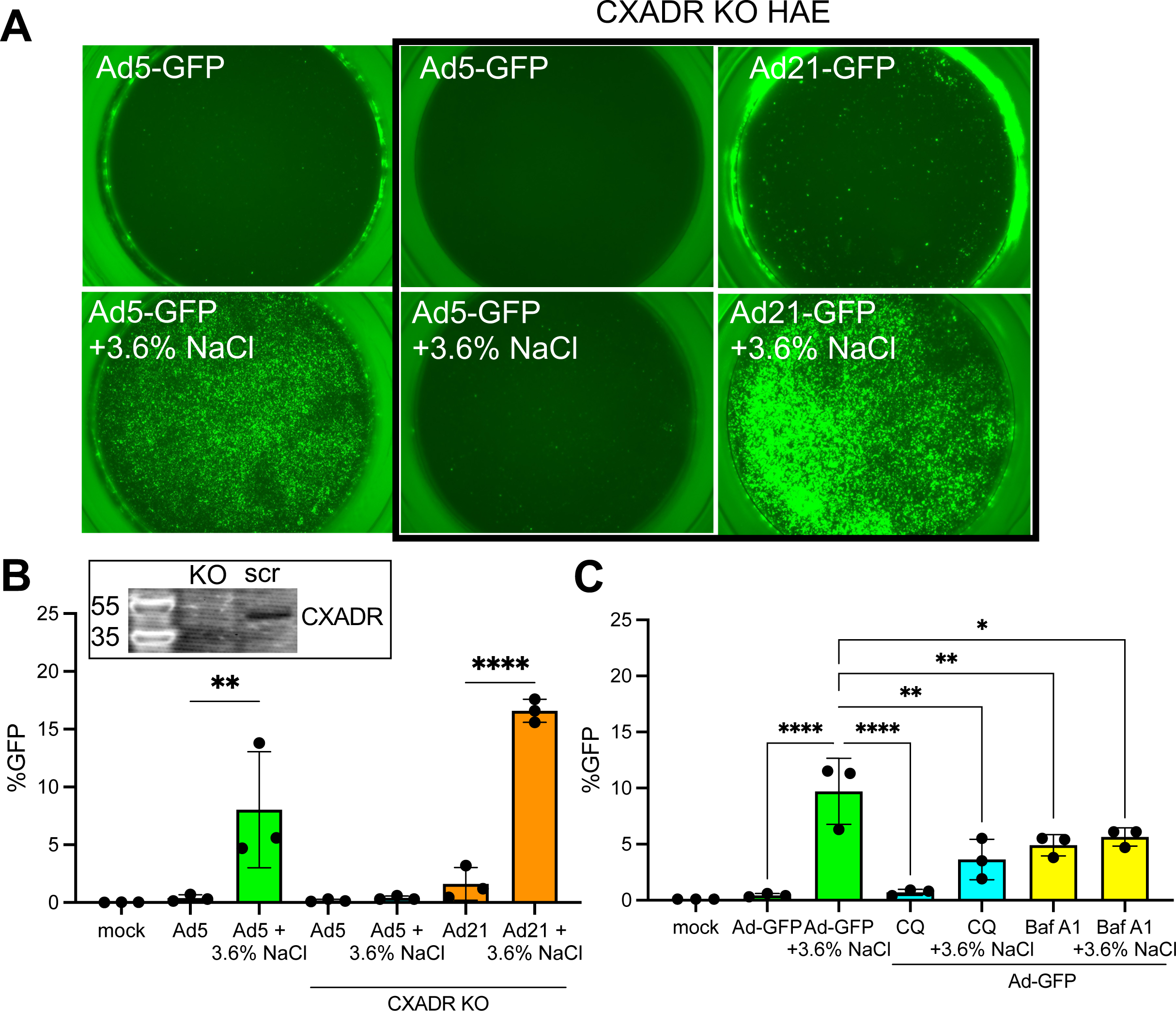
NaCl-mediated transduction requires receptor and low pH entry step. Ad5 utilizes CXADR for entry and Ad21 utilizes CD46. (**A**) CRISPR/Cas9 with gRNA targeted to CXADR was electroporated into HAE and allowed to re-differentiate at air liquid interface for 3 weeks. In parallel with non-electroporated cells from the same donor, Ad5-GFP and Ad21-GFP were applied apically to HAE with or without NaCl for 2 hours. Cultures were imaged 5 days post-transduction and representative images are shown. (**B**) GFP was quantified by flow cytometry. Inset: Western blot for CXADR in parallel on CXADR KO or scrambled airway epithelial cultures. (**C**) HAE were pre-treated with endosomal acidification inhibitors chloroquine (CQ) (200 µM) or bafilomycin A1 (1 µM) for 2 hours. Next, Ad5-GFP (MOI=250) was delivered with or without NaCl (3.6%) for 2 hours. GFP expression was quantified by flow 5 days post-transduction. N=3, *p<0.05, **p<0.005, ****p<0.00005. Statistical differences were determined by one-way ANOVA.

Inhibiting endosomal acidification with chloroquine (CQ) or bafilomycin A1 (Baf A1) significantly decreased the NaCl enhanced transduction (**Fig. 4C**). These results indicate NaCl enhanced transduction requires receptor engagement and low pH endosomal release.

### NaCl formulations enhance lentiviral and AAV gene transfer

We next asked if formulating lentiviral or adeno-associated viral (AAV) vectors with increasing concentrations of NaCl enhanced transduction. An HIV-based viral vector pseudotyped with the baculoviral GP64 envelope glycoprotein^20^ was applied to the apical surface of HAE with 3.5, 4, or 4.5% NaCl. A dose-dependent increase in GFP expression was observed in non-CF HAE (**Fig. 5A**) and we therefore selected 4.5% NaCl for AAV and lentiviral studies. Primary human CF cultures that received GP64 HIV-GFP (MOI=50) also showed remarkable levels of transduction (**Fig. 5B**). Notably, in the presence of NaCl (4.5%), GP64 HIV-CFTR restored the CFTR-dependent anion transport in primary CF HAE. Without NaCl formulation, improvements in CFTR-dependent currents were not observed (**Fig. 5C, D**). Interestingly, NaCl-mediated entry did not enhance transduction of all pseudotyped lentiviral vectors. A screen of additional envelopes from Vesicular Stomatitis Virus (VSVG)^21^, Jaagsiekte Sheep Retrovirus (JSRV)^22^, and Baboon Endogenous Virus (BaEV)^23^ in the presence or absence of NaCl (4.5%) revealed that only GP64 and VSVG showed enhanced transduction (**Supplemental Fig. 4**).

**Fig. 5.**
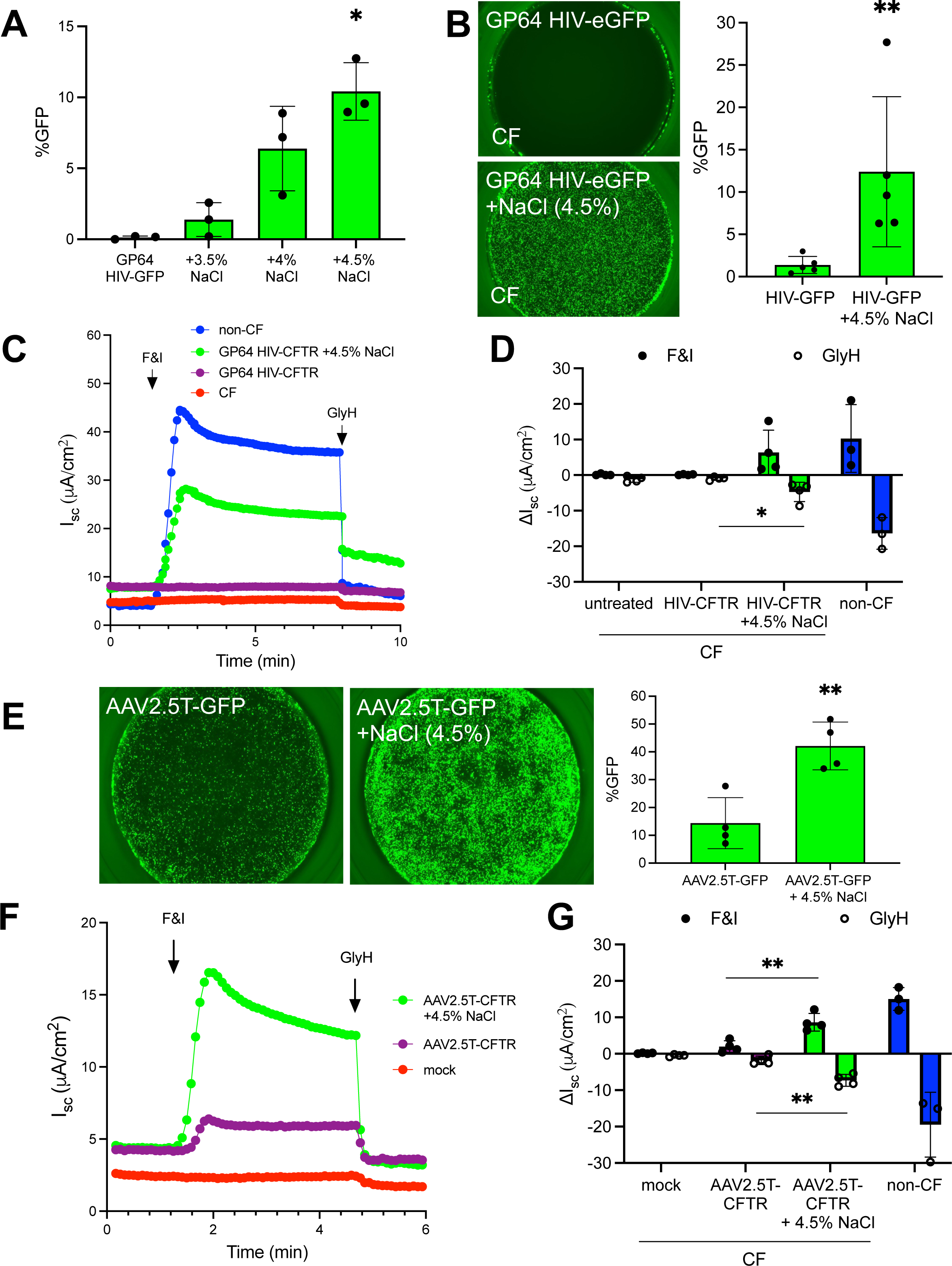
Increasing saline tonicity enhances GP64 pseudotyped lentiviral and AAV transduction. (**A**) GP64 pseudotyped HIV-CMV-eGFP (MOI=50) was co-delivered to airway epithelia with DMEM, 3.5%, 4%, or 4.5% NaCl for 2 hours and then removed. 1 week post-transduction GFP expression was quantified by flow cytometry. (**B**) Parallel CF cultures were transduced with GP64 HIV-CMV-eGFP. GFP expression was visualized by fluorescence microscopy and quantified by flow cytometry 5 days later. (**C, D**) As described in Figure 2, CF airway epithelia were transduced with GP64-HIV-CFTR (MOI=50) formulated with DMEM or 4.5% NaCl for 2 hours and incubated for 1 week. Short circuit current of F&I and GlyH in untreated or treated CF cells compared to non-CF. (**E**) AAV with the 2.5T capsid expressing CMV-eGFP was co-delivered (MOI=100,000) with DMEM or 4.5% NaCl for 2 hours. Airway cells were pre-treated with doxorubicin (1 µM) for 24 hours prior to transduction. 2 weeks post-transduction, airway cultures were imaged and GFP was quantified by flow cytometry. (**F, G**) CF airway cultures were treated overnight with doxorubicin (1 µM) and the next day cells were treated with AAV2.5T-CFTR alone or with 4.5% NaCl for 2 hours. Two weeks post-transduction, electrical properties were measured by Ussing chambers as described in (C, D). N=3, *p<0.05, **p<0.005. Statistical differences were determined by Student’s t-test or one-way ANOVA.

To test the effect of NaCl formulation on AAV-based vectors, we selected the AAV2.5T capsid based on previously established airway tropism^24^. Similar to Ad and lentiviral studies, we applied AAV-GFP (MOI=100,000 vg/cell) to the apical surface of HAE for 2 hours with or without NaCl (4.5%). Doxorubicin (1 µM) was added to the basolateral media 24 hours prior to the 2 hour AAV application. A 2 hour application was chosen to be consistent with the Ad and HIV protocols, however we and others have shown that overnight applications of AAV are effective in the absence of NaCl^25^. Microscopy, flow cytometry quantification, and Ussing chamber assays were performed 2 weeks post-transduction. The epithelia treated with NaCl (4.5%) formulated AAV-GFP achieved significantly higher GFP expression (**Fig. 5E**) and ncreased anion channel activity in responses to F&I and GlyH (**Fig. 5F, G**). These results confirm that ransduction with NaCl formulation is enhanced across all three viral vector systems.

### Increased saline tonicity enhances gene transfer *ex vivo* and *in vivo*

We next asked if formulating vectors in NaCl (5%) would enhance gene transfer in an *ex vivo* model. Using freshly excised newborn pig tracheal explants maintained on Surgifoam, we applied Ad-GFP or AAVH22-GFP (a pig-tropic AAV capsid)^26^ ± NaCl (5%) to the apical surface for 2 hours. One week later, tissues were fixed and imaged by confocal microscopy. Fluorescence was quantified by transduction area per high power field (**Supplemental Fig. 5A, B**). Transduction with both Ad and AAVH22 was markedly enhanced in *ex vivo* tracheal explants when formulated with NaCl (5%).

Improving viral vector transduction *in vivo* has broad implications for advancing gene therapy for lung diseases. We first asked if vectors formulated with NaCl (7%) improved gene transfer to mouse lungs. Ad-uciferase was formulated with NaCl (7%) or NaCl (0.9%) and delivered intratracheally. 5 days post-delivery, animals treated with vectors formulated in 7% NaCl demonstrated significantly increased luciferase expression relative to saline control mice (**Fig. 6A**). We next asked whether NaCl formulation would mprove gene transfer in a large animal model. Newborn pigs received Ad-GFP via intratracheal delivery with PBS or NaCl (5%). Five days after delivery, lungs were collected and divided into 20 regions for systematic quantification as previously described^2^ (**Fig. 6B**). GFP expression was scored using a fluorescence dissecting microscope (0 = no expression, 1 = low expression, 2 = moderate expression, and 3 = high expression). Formulation of Ad-GFP with NaCl (5%) resulted in a significantly increased the GFP+ score compared to PBS (**Fig. 6C-E**). **Fig. 6F** shows representative images of NaCl (5%) treated lung sections at 2x and 10x, respectively. We also tested GP64 HIV-GFP formulated with NaCl (5%) or PBS in pigs. We observed significantly enhanced transduction using NaCl formulation over PBS (**Fig. 6G-I**). In summary, formulating either Ad or lentiviral vectors in 5-7% NaCl enhanced airway and alveolar gene transfer in mice and pigs.

**Fig. 6.**
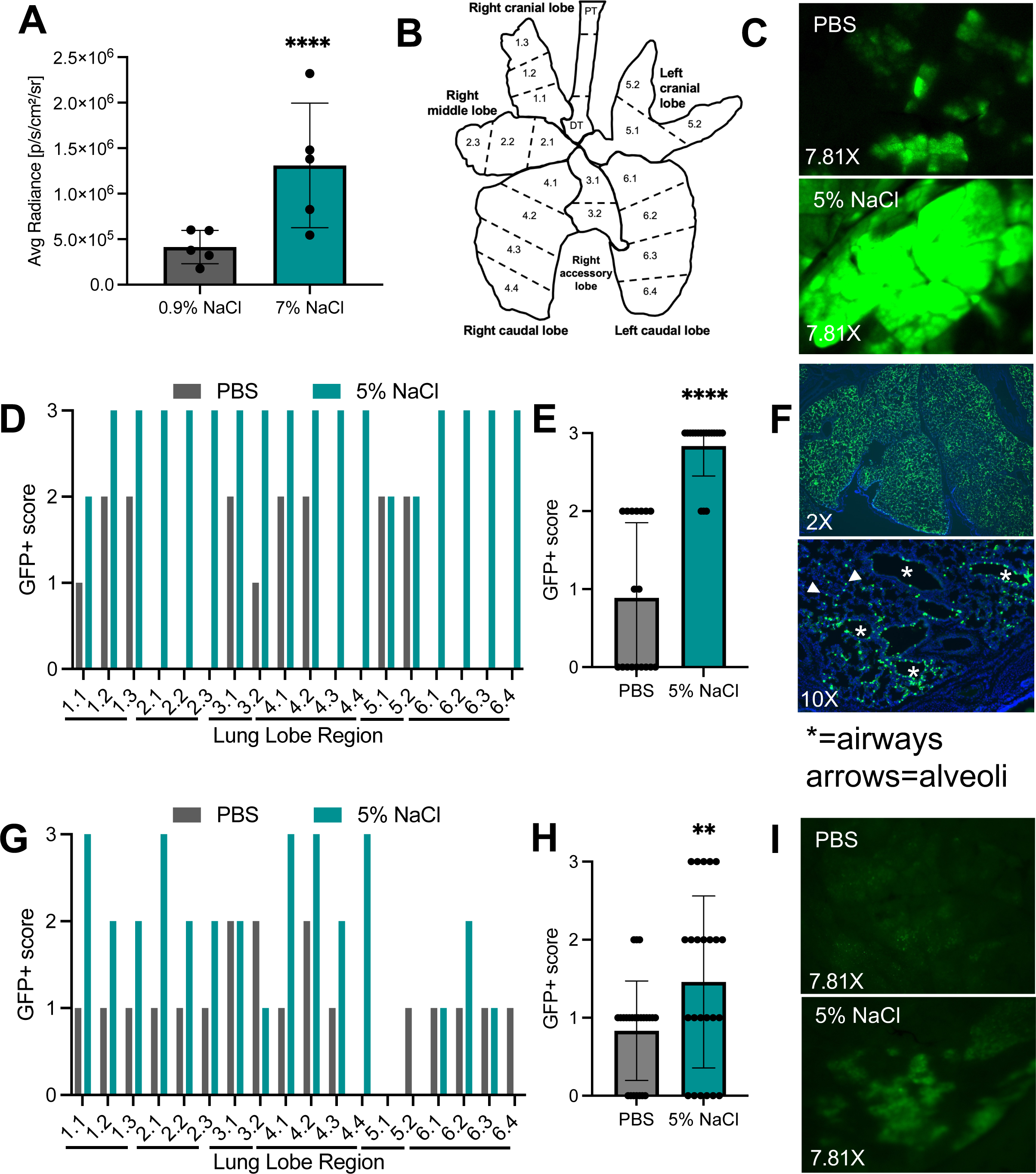
Increased saline tonicity enhances transduction of mouse and pig airways. (**A**) 6-8 week old Balb/c mice received intratracheal Ad-luciferase formulated with NaCl (0.9% or 7% final concentration). Mice were imaged by IVIS 5 days after delivery. N=5. (**B-G**) Ad-GFP was formulated with PBS or 10% NaCl (5% final concentration) and aerosolized intratracheally to newborn pigs. 5 days after delivery lungs were divided into 20 regions (**B**) as previously reported ^2^. (**C**) Representative fluorescent dissecting scope images of GFP expression. (**D**) Individual lung regions were scored using a dissecting fluorescent microscope (0= no expression, 1= low expression, 2=moderate expression, and 3= high expression). (**E**) Combined GFP+ scores. (**F**) Lungs were fixed, processed, sectioned, and counterstained with DAPI. Representative 2X and 10X images from 5% NaCl condition shown. (**G-I**) GP64 HIV-GFP formulated with PBS or NaCl (5% final concentration) was aerosolized to newborn pigs intratracheally. One week later, lungs were collected as described in B. Lobe regions were scored for GFP expression and representative images of fluorescent dissecting microscope shown. Pig experiments: n=1 animal per condition. 10 slides per lung were evaluated. **p<0.005. ****p<0.00005. Statistical differences were determined by Student’s t-test.

## DISCUSSION

Here we show that NaCl (3.6-7%) enhanced the transduction efficiency of Ad, AAV, and lentiviral vectors. This improved transduction restored functional CFTR anion transport in CF airway epithelia across all 3 vector platforms. For adenovirus vectors, the NaCl formulation enhanced transduction remained receptor dependent, requiring a low pH endosomal escape. These *in vitro* findings of enhanced transduction also ranslated to two animal models where formulating Ad or lentiviral vectors with 5% or 7% NaCl increased ransgene expression. Given that 3-7% NaCl is routinely aerosolized in people with CF^27^, these findings have mportant implications for improving viral vector transduction for CF and other lung diseases. Formulating viral vectors with 3.6-7% NaCl could boost therapeutic efficacy, lower the therapeutic index, and overcome previous expression threshold limitations for pulmonary gene therapy.

Formulations including LPC^5^, EGTA^6, 7^, perfluorocarbon^9^, and methylcellulose^8^ have been instrumental in *in vitro* and pre-clinical *in vivo* studies to improve gene transfer efficiency. Interestingly, the discovery that Ad-based viral vectors use a basolateral receptor and gene transfer is boosted by tight junction disruption came to light after the first CF gene therapy clinical trials^4^. Since none of the above mentioned formulations are currently FDA approved for lung administration, each reagent would require a separate evaluation for co-administration with a gene therapy vector. CF gene therapy clinical trials performed to date include vehicles with a 150 mM NaCl concentration, which is equivalent to 0.9% isotonic NaCl (reviewed in^4^). Viral vectors ncluding AAV and lentiviral platforms are under investigation for CF gene therapy applications. 4D Molecular Therapeutics began a Phase I AAV clinical trial in 2022 (clinicaltrials.gov) and Spirovant and Carbon Biosciences are generating preclinical data to pursue clinical trials with AAV and bocavirus vectors. Historically, achieving sufficient gene transfer for phenotypic correction has been challenging^4^. Efforts to overcome these challenges include directed evolution for capsid^24^ and envelope design^20^, cassette modification including promoter^28^, cDNA^29^, and polyadenylation signal, codon optimization^21^, and mmunomodulation (reviewed in^4^). Using a formulation generally recognized as safe for use in people with CF such as NaCl (3-7%) could eliminate a hurdle for approval and advancement of a gene therapy reagent.

The effects of increased saline tonicity formulation on viral vector transduction have not been previously nvestigated. To our knowledge, only one relavant report suggests that increased saline tonicity does not enhance SARS-CoV-2 viral entry^30^. In screening several lentiviral envelope pseudotypes (Supplemental Fig. 4) we observed that vectors with the GP64 and VSVG envelopes benefited from NaCl (4.5%) formulation, while JSRV or BaEV did not. JSRV uses the GPI-linked apical receptor Hyal2^31^ while the BaEV uses SLC1A4 and SLC1A5 as receptors. Although these receptors are present in HAE, transduction was not enhanced with NaCl formulation. These results suggest that entry pathways may influence the effectiveness of NaCl formulations. Indeed, different viruses have differing entry routes, including receptor-mediated endocytosis or fusion at the plasma membrane. Further, it may be that some receptors, co-receptors, and other binding or entry factors may be affected by the presence of high NaCl while others are not. We speculate that virus like particles (VLPs)^32^ pseudotyped with envelopes such as VSVG or GP64 would also be enhanced when formulated with hypertonic saline.

We originally suspected the mechanism for NaCl-mediated enhanced transduction was due to the osmotic strength of the formulation. We know that the airway surface liquid volume is regulated in part by water movement through the cell and that osmotically driven water permeability is regulated by aquaporins to maintain airway epithelial cell homeostasis^33, 34^. When a hypertonic solution is applied to epithelia, an electrochemical gradient generates a driving force for fluid movement across the cell membrane, causing hypertonic shock, followed by an adaptive regulatory increase in cell volume. The non-ionic osmolyte mannitol did not enhance transduction. Instead, we learned that ionic strength is a key determinant in the mechanism by which NaCl formulation enhance vector transduction. We speculate that NaCl formulation may enhance transduction, in part, by increasing receptor access by disrupting tight junctions or by enhancing endosomal escape. We observed that NaCl, KCl, and the charged sugar N-methyl-D-glucamine enhanced transduction in a dose-dependent manner, although NaCl had the greatest impact in a given percent solution. These findings suggest that Na^+^ regulation of the airway extracellular environment has the greatest mpact on viral transduction^35^.

As mentioned in the Results, a concentration of 3.6% final concentration of NaCl was serendipitously chosen for most of the studies using Ad because it was initially diluted 1:1 from a 7.2% pharmaceutical grade stock solution. For *in vitro* studies in HAE, our dose response results suggest that 4.5% NaCl may be the best compromise between efficacy and ultimate toxicity at doses of NaCl >6%. This was true for all vectors ested. Of note, attempts to improve gene transfer of cultured cells on plastic using hypertonic saline resulted n lifting of the cells. Our pilot studies *in vivo* suggested that vector formulations of 5-7% NaCl had the greatest impact on gene transfer to the lungs of mice and pigs. This suggests that the dose limitation *in vitro* may not apply to *in vivo* gene transfer and higher initial concentrations of NaCl may be necessary to counteract the immediate dilution of the encountered airway surface liquid.

The goal of this study was to investigate whether FDA approved agents could improve lung gene therapy efficacy. Hypertonic saline is a routinely used therapy for people with CF and other lung conditions and could be rapidly adopted in vector formulations. We show evidence that formulating Ad, AAV or lentiviral vectors with NaCl could advance gene therapy development for CF. Importantly, this finding has broad mplications for other genetic lung diseases such as alveolar type 2 deficiencies (ABCA3 Deficiency, Surfactant Protein B Deficiency) or Primary Ciliary Dyskenesia.

## MATERIALS AND METHODS

### Ethics statement

Primary airway epithelia from human CF and non-CF donors were isolated from discarded tissue, autopsy, or surgical specimens. Cells were provided by The University of Iowa *In Vitro* Models and Cell Culture Core Repository. Information that could be used to identify a subject was not provided. All studies involving human subjects received University of Iowa Institutional Review Board approval (Protocol #230167). Mice and pig experimental protocols were reviewed and approved by the University of Iowa Institutional Animal Care and Use Committee (IACUC), in accordance with the United States Department of Agriculture and National Institutes of Health guidelines.

### Human airway epithelial cells

The University of Iowa *In Vitro* Models and Cell Culture Core cultured and maintained HAE as previously described^36^. Briefly, following enzymatic disassociation of trachea and bronchus epithelia, the cells were seeded onto collagen-coated, polycarbonate Transwell inserts (0.4 μm pore size; surface area = 0.33 cm^2^; Corning Costar, Cambridge, MA). HAE were submerged in Ultroser G (USG) medium for 24 hours (37°C and 5% CO_2_) at which point the apical media is removed to encourage polarization and differentiation at an air-liquid interface. Transepithelial electrical resistance was measured using an Ohmmeter (Ω·μm^2^).

### Viral vector production and formulation

Vectors were produced by the University of Iowa Viral Vector Core (https://medicine.uiowa.edu/vectorcore/). Ad5 and Ad21 CMV-eGFP CMV-mCherry, or F5Tg83-CFTR was produce, purified, and titered as previously described^37^. AAV2/2.5T, AAV2/H22 CMV-eGFP or F5Tg83-CFTRι1R was produced by triple transfection. Lentiviral HIV-CMV-eGFP or PGK-CFTR vectors pseudotyped with GP64 were produced by a four-plasmid transfection method as previously described^38, 39^ and titered using droplet digital PCR ^40^ and/or by flow cytometry. Similarly, VSVG, JSRV and BaEV pseudotyped lentiviral vectors were made in-house by four-plasmid transfection and titered by flow cytometry. Viral vectors were formulated with indicated NaCl concentrations (presented as final concentrations). The following MOIs were used for each vector: Ad=250, AAV=100,000, and HIV=50. Ad studies were performed by mixing pharmaceutical grade 7.2% NaCl with Ad-GFP (3.6% final). Additional concentrations were made from a 10% NaCl solution of Sodium Chloride (Research Products International, Mount Prospect, IL) in UltraPure Distilled Water (Invitrogen, Waltham, MA). Vectors were formulated with NaCl in a final volume of 50 µl and applied to the apical surface of HAE for 2 hours. AAV transduced cultures were pre-treated with indicated concentraions of doxorubicin [1 µM] for 24 hours prior to 2 hour vector application. Other formulations tested include lysophosphatidylcholine (LPC) (9008-30-4, Millipore Sigma, St. Louis, MO), Ethylene glycol-bis-2-aminoethylether)-N,N,N’N’-tetraacetic acid (EGTA) (Research Products International, Mount Prospect, IL), and surfactant (Infasurf, pharmaceutical grade).

### Fluorescence microscopy and flow cytometry

GFP images were acquired using a Keyence All-in-one Fluorescence Microscope BZ-X series (Osaka, Japan). 0.33 cm^2^ transwells were imaged at 2X magnification. GFP expression was quantified by flow cytometry as previously reported^39, 41^. Briefly, cells were stained with a fixable LIVE/DEAD stain (Thermo Fisher Scientific, Waltham, MA), lifted in Accutase at 37°C for 30 minutes, and run through an Attune NxT Flow Cytometer (Thermo Fisher Scientific, Waltham, MA). Cells were treated with the Foxp3/Transcription Factor Staining Buffer Set (Thermo Fisher Scientific, Waltham, MA, USA) according to manufacturer’s recommendations and stained for 1 hour at 4°C with the following antibodies: NGFR (345110; 1:600, BioLegend, San Diego, CA, USA), α-tubulin (NB100-69AF405, 1:300, Novus, Centennial, CO, USA) and CD66c (12-0667-42, 1:600, Invitrogen, Waltham, MA). Expression was gated on live cells.

### LDH release assay

LDH release was quantified according to the manufacturer’s recommendations (LDH-Glo Cytotoxicity Assay, Promega, Madison, WI). Briefly, basolateral media from each condition was collected in a 96 well plate. The LDH detection reagents were mixed in a 1:1 ratio and applied to each well and incubated for 30 minutes. Luminescence was recorded using a SpectraMax i3x plate reader.

### Dextran permeability assay

Tight junction permeability was assessed by passage of 3000-5000 average molecular weight Fluorescein sothiocyanate-dextran (FD4, Sigma-Aldrich, St. Louis, MO) from the apical to basolateral side of well-differentiated human airway epithelia. Carbonyl cyanide 3-chlorophenylhydrazone (CCCP) (Sigma-Aldrich, St. Louis, MO) serves as a positive control for membrane permeabilization and was pre-treated (50 µM) overnight prior to addition of dextran. 250 mM EGTA pre-treatment for 2 hours was used for a positive experimental tight junction disruption control. Cells were left untreated or pre-treated with NaCl ranging from 1-7% for 2 hours. At the time of the assay, pre-treatments were removed and fresh basolateral media was added. 100 µl of 1 mg/ml dextran was applied apically for 30 minutes at 37°C. Basolateral media was assessed for fluorescent units using a SpectraMax i3x plate reader.

### SEM

Scanning electron microscopy (SEM) samples were processed by the Johns Hopkins Institute for Biomedical Sciences Microscope Facility as a fee for service (https://microscopy.jhmi.edu/Services/EMCostStruct.html).

### Electrophysiology

Short circuit current (I_sc_) and conductance (G_sc_) was measured as previously described^39^. Briefly, cells were pre-stimulated with forskolin and IBMX (F&I) overnight and their bioelectric properties were quantified by Ussing chamber analysis. Assay protocol is as follows: amiloride, 4,4′-diisothiocyano-2,2′-stilbenedisulfonic acid (DIDS), F&I, and GlyH-101 (GlyH). Results are reported as change in short circuit current (ΔI_sc_) and conductance (ΔG_sc_) in response to F&I or GlyH.

### Western blot

Protein from human airway epithelia transduced with Ad-CFTR alone or Ad-CFTR+NaCl (3.6%) was harvested using RIPA buffer. 20 µg of each sample was loaded on a Criterion Tris-Glycine 4-20% gel and run at 110V for 1 hour and transferred overnight onto a PVDF membrane at 110 µA at 4°C. After membrane blocking, each membrane was probed for using the following primary antibodies: mouse anti-CFTR 596 (1:1000) (UNC) or CAR anti-rabbit (CXADR) (1:100) (PA5-31175, Invitrogen, Waltham, MA) was applied for 2 hours. Following TBST washes, the LiCor secondary antibody was used at 1:10,000 for 30 minutes. The membrane was imaged using an Odyssey imager.

### Sugar tonicity and ionic vs. non-ionic osmolytes

KCl, mannitol, and N-methyl-D-glucamine (NMDG) gluconate were formulated 1-7% and co-delivered with Ad-GFP (MOI=250) for 2 hours apically. Molarities and osmolarities are presented in Table 1. Sodium chloride (NaCl), mannitol, sodium gluconate, and potassium chloride (KCl) were made with equal percentages (3.6%) or equal molarity (1,23 mOsmol/L) to NaCl as shown in Table 2. Each solution was mixed with Ad-GFP (MOI=250) and applied apically to airway epithelia for 2 hours. GFP expression was quantified by flow cytometry 5 days post-transduction.

### CXADR KO

CRISPR/Cas9 was used to knockout the coxsackie and adenovirus receptor CXADR by nucleofection (Lonza, Basel Switzerland). Well-differentiated human airway epithelia were lifted in TrypLE and electroporated with ribonucleoproteins (RNP). CXADR Exon 4 was targeted with the following guide RNAs: gRNA 1: ACGTAACATCTCGCACCTGAAGG and gRNA 2: AGTACCTGCTAACCATGAAGTGG ^17^. Alt-R S.p. HiFi Cas9 nuclease (1081058, IDT, Coralville, IA) was mixed with crRNA and trcrRNA duplex to form the RNP. RNP and cells were mixed with the nucleofector solution and electroporated using program U-024 and seeded in a 6 well plate in Pneumacult ExPlus media. 2 days later, cells were lifted and reseeded on a Corning 3413 collagen coated membranes. Cells were maintained in Pneumacult ALI-M media for 3 weeks to differentiate. Ad5-GFP (CXADR-dependent) and Ad21-GFP (CXADR-independent) were formulated with DMEM or NaCl (3.6%) and applied to the apical surface of airway epithelia for 2 hours. GFP expression was quantified 5 days post-transduction by flow cytometry.

### Inhibitors of endosomal acidification

Endosomal acidification inhibitors choloroquine (CQ) (200 µM) (C6628, Millipore Sigma, St. Louis, MO) and bafilomycin A1 (Baf A1) (1 µM) (19148, Millipore Sigma, St. Louis, MO) were used to pre-treat airway epithelia for 2 hours prior to Ad-GFP transduction in the presence or absence of NaCl (3.6%). GFP was quantified by flow cytometry 5 days post-transduction.

### Pig explants transduction

0.25 cm^2^ tracheal explants from newborn pigs were cultured on Surgifoam (1972, Ethicon, Raritan, NJ) for 3-5 days prior to transduction. Ad5-mcherry (1x10^9^ TU), AAVH22-GFP (2x10^10^ vg) were applied with or without NaCl (5%) to the apical surface of the trachea by inverting the trachea onto a tissue culture dish containing the viral solution for 2 hours then returned to an upright position on Surgifoam. 5 days post-ransduction, tracheal explants were mounted on a microscope slide and imaged for fluorescent expression using confocal microscopy. Expression was quantified using Image J (FIJI) by measuring fluorescence ntensity per high power field.

### Adenoviral transduction of mouse airways

6-8 week old Balb/c mice were sedated with ketamine/xylazine (87.5 + 2.5 mg/kg). Ad-CMV-firefly uciferase was delivered intratracheally with 1x10^9^ TU formulated with NaCl (0.9% or 7% final). 5 days later mice were given intraperitoneal D-luciferin (50227, Millipore Sigma, St. Louis, MO) imaged for uminescence expression using the Xenogen IVIS-200.

### Viral gene transfer in pig airways

Newborn pigs were sedated using isoflurane and viral vector formulated with NaCl (0.9% or 5% final) and aerosolized intratracheally using a MADgic Laryngo-Tracheal Mucosal Atomization Device (Teleflex, Morrisville, NC). Doses for each vector for each pig were as follows: 2.8x10^10^ infectious genomic units (IGU) of helper-dependent Ad-CMV-GFP and 7x10^8^ TU of GP64 HIV-CMV-GFP. 1 week later, pigs were humanely euthanized and lungs were analyzed for GFP expression. Lungs were systematically divided and fixed in 4% paraformaldehyde, subjected to a sucrose gradient, and embedded in OCT for cryosectioning. Sectioned lung slides were mounted using DAPI and imaged using a Keyence All-in-one Fluorescence Microscope BZ-X series (Osaka, Japan).

### Statistics

Student’s two-tailed t-test, one-way analysis of variance (ANOVA) with Tukey’s multiple comparison test, or two-way ANOVA with Dunnett’s multiple comparison test were used to analyze differences in mean values between groups. Results are expressed as mean ± SEM. P values ≤ 0.05 were considered significant. R^2^ values in Fig. 1E represent exponential growth of nonlinear regression.

## Acknowledgments

We thank Ian Thornell for the insightful conversations which helped us identify ionic strength as the mechanism for NaCl-mediated vector transduction. We thank Amber Vu for her work on lentiviral envelopes and Soumba Traore for guidance on CXADR knockout of airway epithelia. We thank Linda Powers, Brie Hilken, and Nick Gansemer for orchestrating the pig experiments and delivering the viral vector. We thank Ian Thornell, Lynda Ostedgaard, Cami Hippee, Lorellin Durnell, and Stephanie Clark for their critical review of this manuscript.

## Funding

National Institutes of Health grant R01HL133089 (PLS)

National Institutes of Health grant P01HL152960 (PBM)

National Institutes of Health grant R01HL171035 (PLS, PBM)

National Institutes of Health grant P01HL51670 (PBM)

Cystic Fibrosis Foundation grant SINN22G0 (PLS)

Cystic Fibrosis Foundation grant SINN23G0 (PLS)

Cystic Fibrosis Foundation grant STOLTZ19 (PLS)

Emily’s Entourage grant (PLS)

The University of Iowa Center for Gene Therapy, National Institutes of Health grant P30DK54759 (PLS)

Roy J. Carver Chair in Pulmonary Research (PBM)

American Society of Gene and Cell Therapy Career Development Award (ALC)

## Author contributions

Conceptualization: ALC, PBM, PLS

Methodology: ALC, LML, KN, CMB

Investigation: ALC

Visualization: ALC, PBM, PLS

Funding acquisition: PBM, PLS

Project administration: ALC

Supervision: PBM, PLS

Writing – original draft: ALC

Writing – review & editing: ALC, PBM, PLS

---

**Supplemental Fig. 1.**
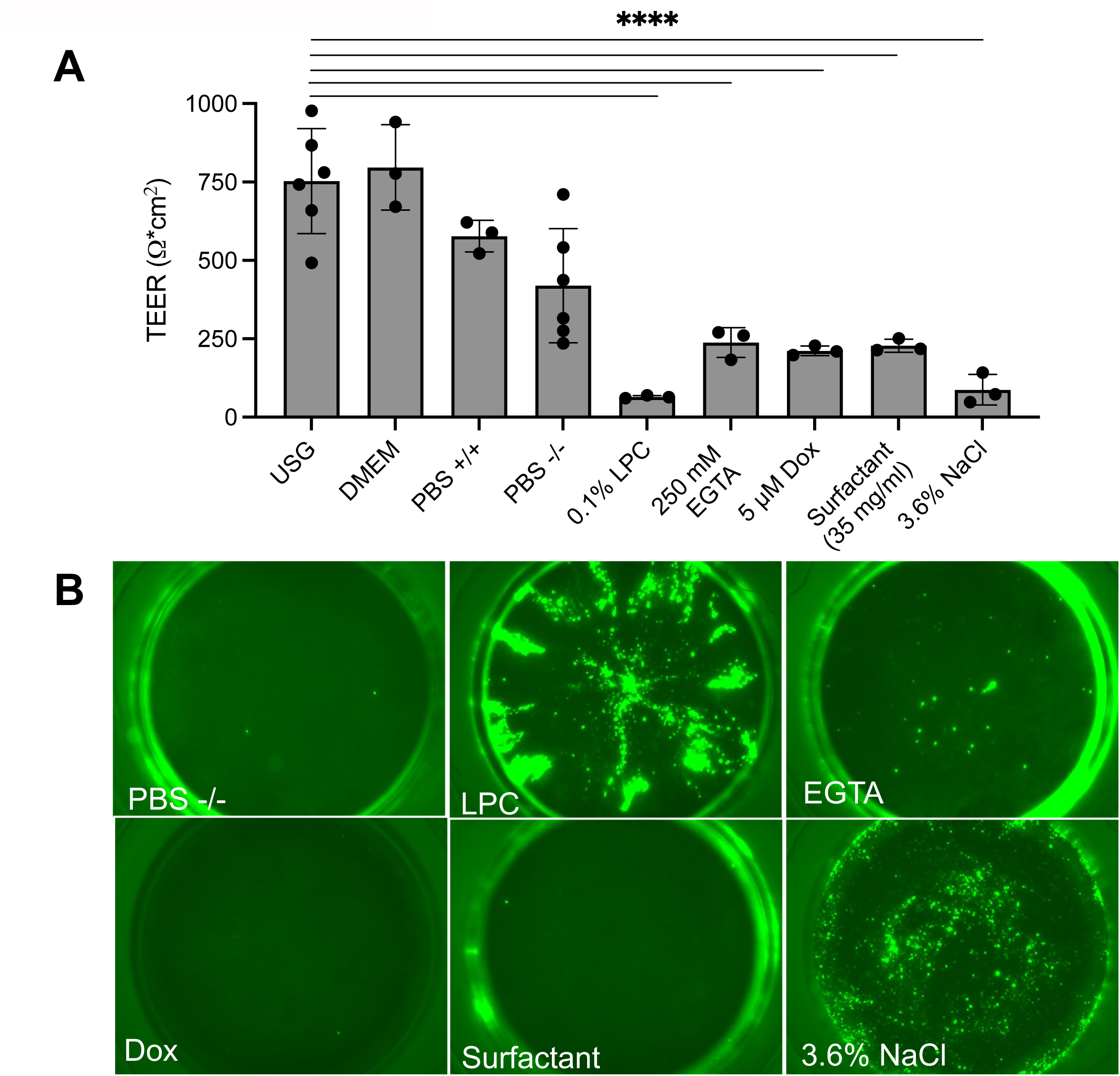
Clinically approved agents decrease transepithelial resistance but only hypertonic saline (3.6% NaCl) enhances gene transfer. (**A**) Transepithelial electrical resistance (TEER) of HAE was measured using an ohmmeter (normalized to 0.33 cm^2^ transwell area) after 5 minute incubation with indicated reagents: Ultraser G (USG), Dulbecco’s Modified Eagle Medium (DMEM), phosphate buffered saline (PBS) with (+/+) or without (-/-) calcium and magnesium, 0.1% lysophosphatidylcholine (LPC), 250 mM EGTA, 5 µM doxorubicin (Dox), surfactant (Infasurf ) (35 mg/ml), or 3.6% NaCl. (B) Ad-GFP (MOI=250) was applied apically to HAE with PBS -/-, LPC, EGTA, Dox, Surfactant, or 3.6% NaCl for 2 hours. Cultures were imaged 5 days post-transduction. ****p<0.00005.

**Supplemental Fig. 2.**
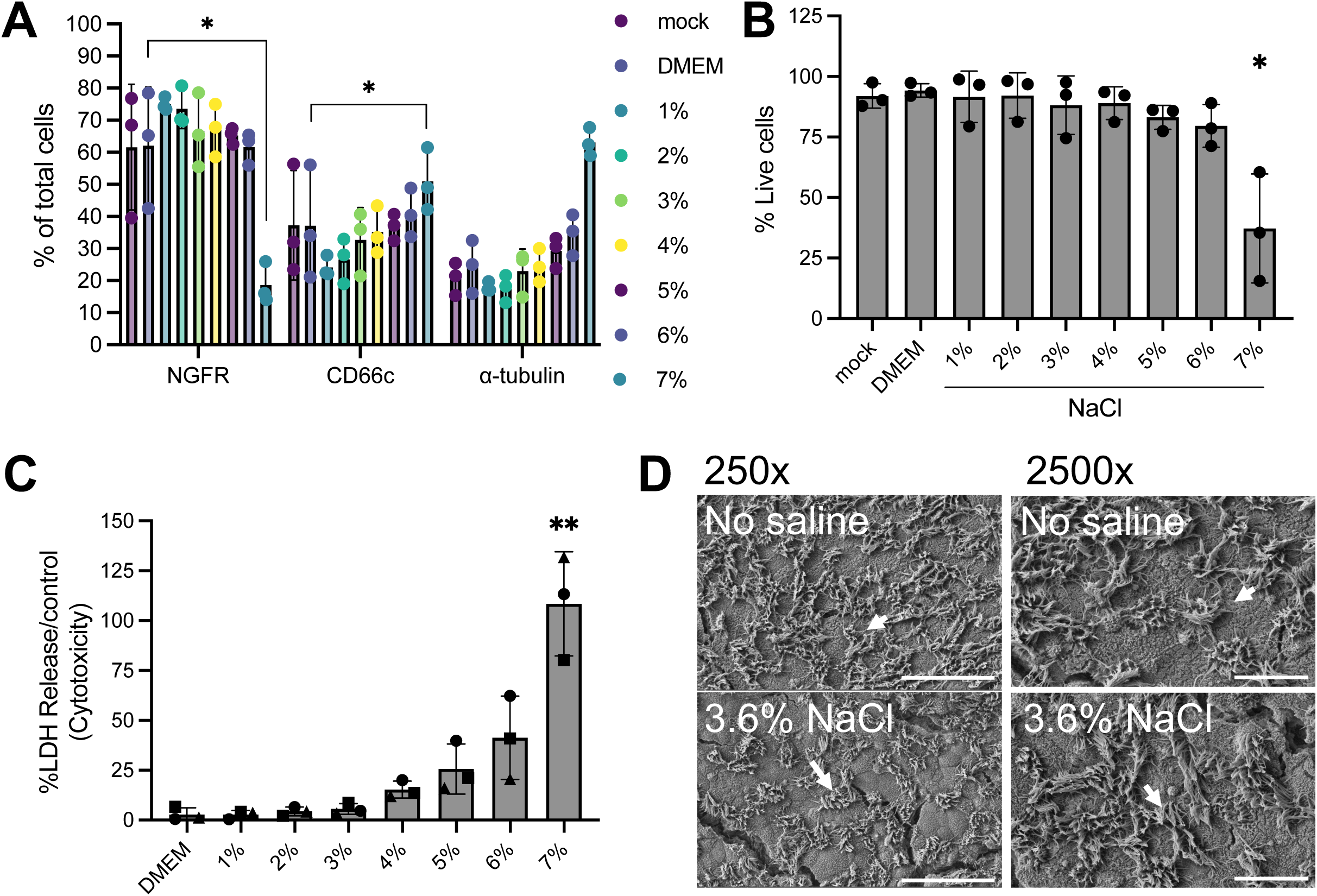
Effects of NaCl treatment on primary human airway epithelia. (**A**) Distribution of cell types with increasing %NaCl 5 days after treatment. Nerve growth factor receptor (NGFR)=basal cells, CD66c=secretory cells, α-tubulin=ciliated cells. (**B**) HAE from (**A**) were treated with LIVE/DEAD followed by immediated fixation prior to assaying by flow cytometry. %Live cells reported for 1-7% NaCl. (**C**) Cytotoxicity as measured by %LDH release for 1-7% NaCl. (**D**) SEM images after 2 hour treatment with DMEM or NaCl (3.6%). N=3, p<0.05. Scale bar=50 µm. Arrows indicate cilia.

**Supplemental Fig. 3.**
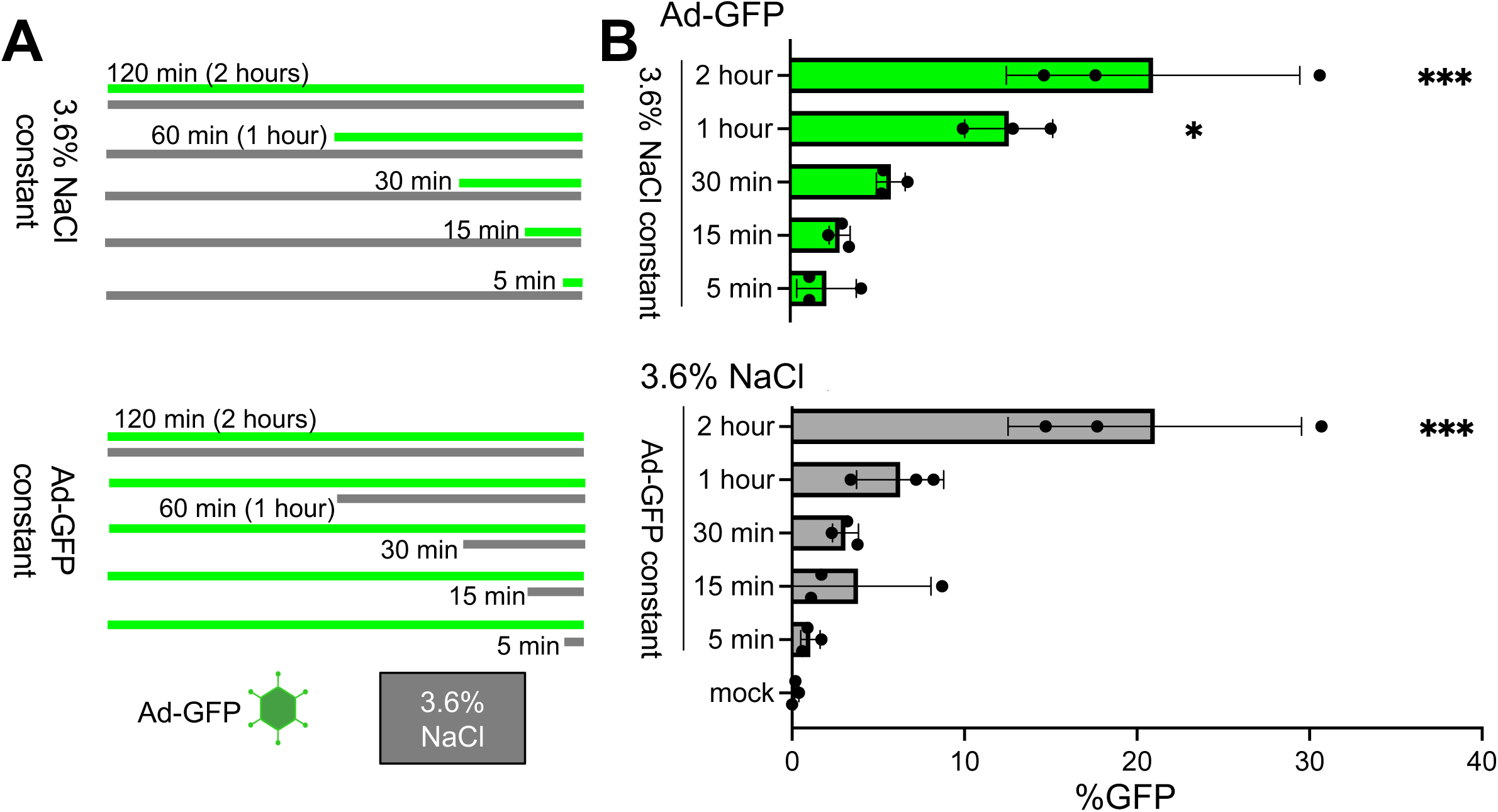
NaCl-mediated entry of Ad-GFP is time dependent. (**A**) Experiment schematic. Ad-GFP (green bar) was applied for 2 hours with 3.6% NaCl (gray bar) present at 120, 60, 30, 15, or 5 minutes. Parallel cultures were incubated with 3.6% NaCl for 2 hours with Ad-GFP present for 120, 60, 30, 15, or 5 minutes. The 2 hour co-treatment group was shared between groups. (**B, C**) Images were acquired 5 days post-transduction. Representative fluorescence microscopy images at each time interval. (**D**) GFP quantification by flow cytometry. N=3, *p<0.05, ***p<0.0005. Statistical differences were determined by one-way ANOVA.

**Supplemental Fig. 4.**
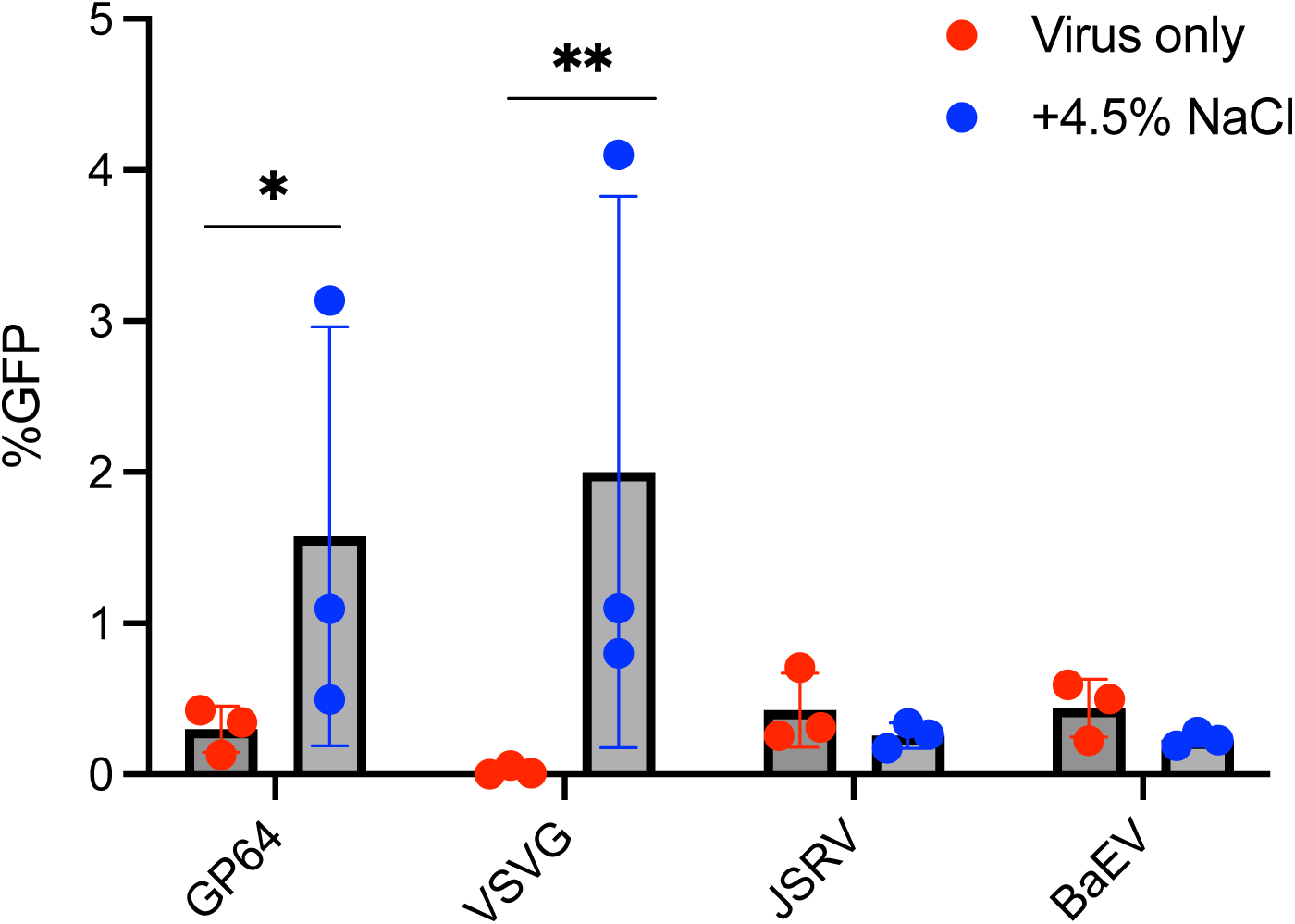
NaCl-mediated entry does not enhance all lentiviral pseudotypes. Baculovirus glycoprotein 64 (GP64), Vesicular Stomatitis Virus (VSVG), Jaagsiekte Sheep Retrovirus (JSRV), Baboon Endogenous Virus (BaEV) pseudotyped HIV-GFP (MOI=10) were treated for 2 hours without and with 4.5% NaCl. GFP expression was quantified 5 days later by flow cytometry. N=3, *p<0.05, **p<0.005.

**Supplemental Fig. 5.**
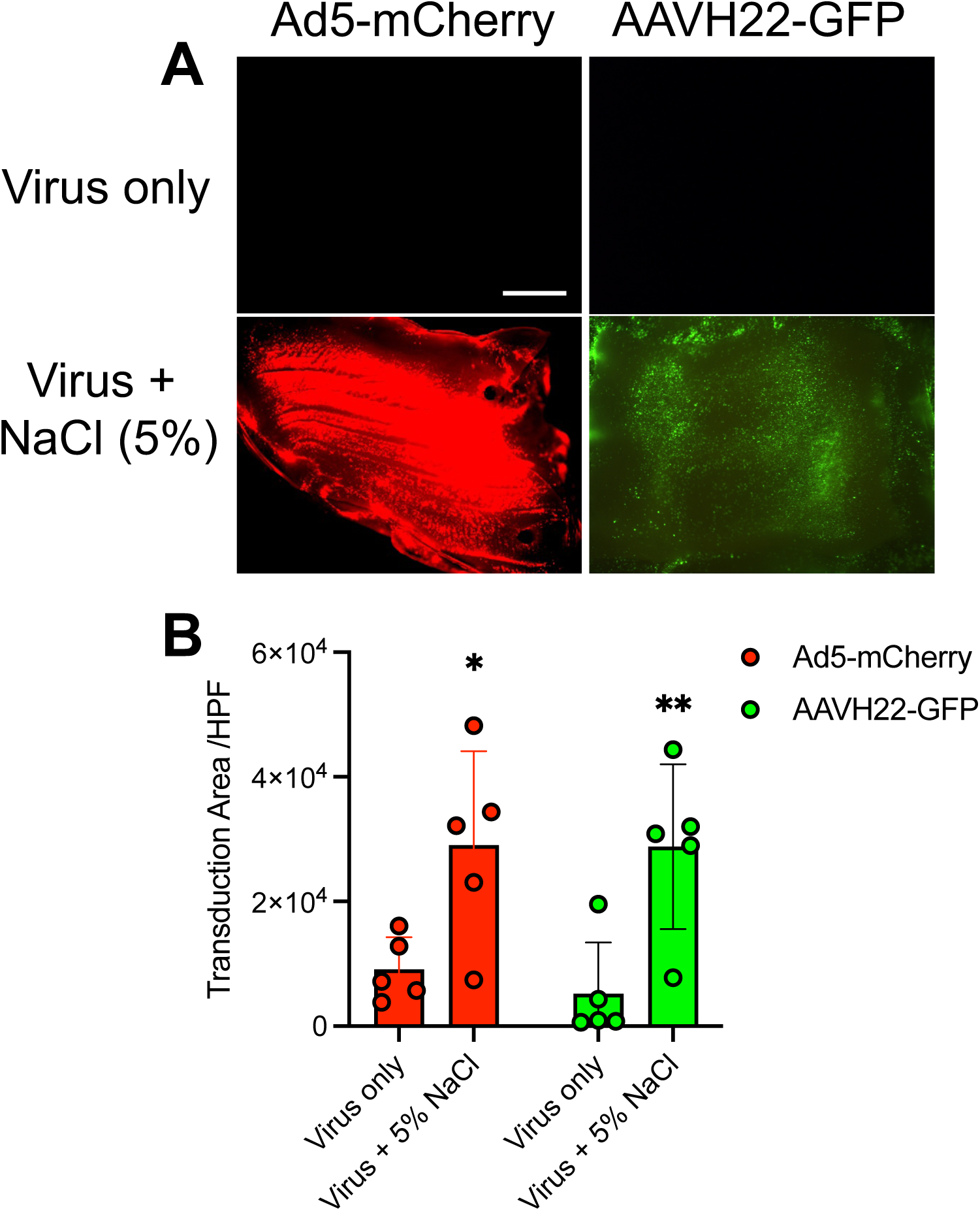
NaCl formulated with Ad and AAV enhances transduction in pig tracheal explants. Each vector (Ad-mCherry 1x10^9^ TU; AAVH22-GFP 2x10^10^ vg) was formulated with NaCl (5% final concentration) and applied to the apical surface of pig tracheal explants for 2 hours. (**A**) mCherry or GFP fluorescence expression was visualized 5 days post-transduction. (**B**) Images were quantified using ImageJ and reported as fluorescent intensity per high power field (HPF). N=5, *p<0.05, **p<0.005.

## REFERENCES

1. Cooney, A.L., et al. Lentiviral-mediated phenotypic correction of cystic fibrosis pigs. JCI Insight 1 (2016).

2. Cooney, A.L. et al. Widespread airway distribution and short-term phenotypic correction of cystic fibrosis pigs following aerosol delivery of piggyBac/adenovirus. Nucleic Acids Res 46, 9591–9600 (2018).

3. Cooney, A.L. et al. A Novel AAV-mediated Gene Delivery System Corrects CFTR Function in Pigs. Am J Respir Cell Mol Biol 61, 747–754 (2019).

4. Cooney, A.L., McCray, P.B., Jr. & Sinn, P.L. Cystic Fibrosis Gene Therapy: Looking Back, Looking Forward. Genes (Basel) 9 (2018).

5. Cmielewski, P., Anson, D.S. & Parsons, D.W. Lysophosphatidylcholine as an adjuvant for lentiviral vector mediated gene transfer to airway epithelium: effect of acyl chain length. Respir Res 11, 84 (2010).

6. Wang, G. et al. Increasing epithelial junction permeability enhances gene transfer to airway epithelia In vivo. Am J Respir Cell Mol Biol 22, 129–138 (2000).

7. Wang, G. et al. Influence of cell polarity on retrovirus-mediated gene transfer to differentiated human airway epithelia. J Virol 72, 9818–9826 (1998).

8. Sinn, P.L., Shah, A.J., Donovan, M.D. & McCray, P.B., Jr. Viscoelastic gel formulations enhance airway epithelial gene transfer with viral vectors. Am J Respir Cell Mol Biol 32, 404–410 (2005).

9. Kazzaz, J.A. et al. Perfluorochemical liquid-adenovirus suspensions enhance gene delivery to the distal lung. Pulm Med 2011, 918036 (2011).

10. Hogman, M., Mork, A.C. & Roomans, G.M. Hypertonic saline increases tight junction permeability in airway epithelium. Eur Respir J 20, 1444–1448 (2002).

11. Elkins, M.R. & Bye, P.T. Mechanisms and applications of hypertonic saline. J R Soc Med 104 **Suppl 1**, S2–5 (2011).

12. Tarran, R., Donaldson, S. & Boucher, R.C. Rationale for hypertonic saline therapy for cystic fibrosis lung disease. Semin Respir Crit Care Med 28, 295–302 (2007).

13. Excoffon, K.J. et al. Isoform-specific regulation and localization of the coxsackie and adenovirus receptor in human airway epithelia. PLoS One 5, e9909 (2010).

14. Terlizzi, V., Masi, E., Francalanci, M., Taccetti, G. & Innocenti, D. Hypertonic saline in people with cystic fibrosis: review of comparative studies and clinical practice. Ital J Pediatr 47, 168 (2021).

15. Cooney, A.L., Singh, B.K. & Sinn, P.L. Hybrid nonviral/viral vector systems for improved piggyBac DNA transposon in vivo delivery. Mol Ther 23, 667–674 (2015).

16. Lei, L. et al. CFTR-rich ionocytes mediate chloride absorption across airway epithelia. J Clin Invest 133 (2023).

17. Ortiz-Zapater, E. et al. Epithelial coxsackievirus adenovirus receptor promotes house dust mite-induced lung inflammation. Nat Commun 13, 6407 (2022).

18. Krishnamurthy, S., et al. Functional correction of CFTR mutations in human airway epithelial cells using adenine base editors. Nucleic Acids Res 49, 10558-10572 (2021).

19. Sirena, D. et al. The human membrane cofactor CD46 is a receptor for species B adenovirus serotype 3. J Virol 78, 4454–4462 (2004).

20. Sinn, P.L. et al. Novel GP64 envelope variants for improved delivery to human airway epithelial cells. Gene Ther 24, 674–679 (2017).

21. Marquez Loza, L.I., et al. Increased CFTR expression and function from an optimized lentiviral vector for cystic fibrosis gene therapy. Mol Ther Methods Clin Dev 21, 94–106 (2021).

22. Sinn, P.L. et al. Gene transfer to respiratory epithelia with lentivirus pseudotyped with Jaagsiekte sheep retrovirus envelope glycoprotein. Hum Gene Ther 16, 479–488 (2005).

23. Girard-Gagnepain, A. et al. Baboon envelope pseudotyped LVs outperform VSV-G-LVs for gene transfer into early-cytokine-stimulated and resting HSCs. Blood 124, 1221–1231 (2014).

24. Excoffon, K.J. et al. Directed evolution of adeno-associated virus to an infectious respiratory virus. Proc Natl Acad Sci U S A 106, 3865–3870 (2009).

25. Zhang, L.N. et al. Dual therapeutic utility of proteasome modulating agents for pharmaco-gene therapy of the cystic fibrosis airway. Mol Ther 10, 990–1002 (2004).

26. 26. Steines, B., et al. CFTR gene transfer with AAV improves early cystic fibrosis pig phenotypes. *JCI Insight* 1, e88728 (2016).

27. Elkins, M.R. et al. A controlled trial of long-term inhaled hypertonic saline in patients with cystic fibrosis. N Engl J Med 354, 229–240 (2006).

28. Yan, Z. et al. Optimization of Recombinant Adeno-Associated Virus-Mediated Expression for Large Transgenes, Using a Synthetic Promoter and Tandem Array Enhancers. Hum Gene Ther 26, 334–346 (2015).

29. Ostedgaard, L.S. et al. Cystic fibrosis transmembrane conductance regulator with a shortened R domain rescues the intestinal phenotype of CFTR-/-mice. Proc Natl Acad Sci U S A 108, 2921–2926 (2011).

30. Machado, R.R.G. et al. Inhibition of Severe Acute Respiratory Syndrome Coronavirus 2 Replication by Hypertonic Saline Solution in Lung and Kidney Epithelial Cells. ACS Pharmacol Transl Sci 4, 1514–1527 (2021).

31. Rai, S.K., DeMartini, J.C. & Miller, A.D. Retrovirus vectors bearing jaagsiekte sheep retrovirus Env transduce human cells by using a new receptor localized to chromosome 3p21.3. J Virol 74, 4698–4704 (2000).

32. Banskota, S. et al. Engineered virus-like particles for efficient in vivo delivery of therapeutic proteins. Cell 185, 250–265 e216 (2022).

33. Kreda, S.M., Gynn, M.C., Fenstermacher, D.A., Boucher, R.C. & Gabriel, S.E. Expression and localization of epithelial aquaporins in the adult human lung. Am J Respir Cell Mol Biol 24, 224–234 (2001).

34. Verkman, A.S., Matthay, M.A. & Song, Y. Aquaporin water channels and lung physiology. Am J Physiol Lung Cell Mol Physiol 278, L867–879 (2000).

35. Bartoszewski, R., Matalon, S. & Collawn, J.F. Ion channels of the lung and their role in disease pathogenesis. Am J Physiol Lung Cell Mol Physiol 313, L859–L872 (2017).

36. Karp, P.H. et al. An in vitro model of differentiated human airway epithelia. Methods for establishing primary cultures. Methods Mol Biol 188, 115–137 (2002).

37. Anderson, R.D., Haskell, R.E., Xia, H., Roessler, B.J. & Davidson, B.L. A simple method for the rapid generation of recombinant adenovirus vectors. Gene Ther 7, 1034–1038 (2000).

38. Sinn, P.L., Coffin, J.E., Ayithan, N., Holt, K.H. & Maury, W. Lentiviral Vectors Pseudotyped with Filoviral Glycoproteins. Methods Mol Biol 1628, 65–78 (2017).

39. Cooney, A.L., Thurman, A.L., McCray, P.B., Jr., Pezzulo, A.A. & Sinn, P.L. Lentiviral vectors transduce lung stem cells without disrupting plasticity. Mol Ther Nucleic Acids 25, 293–301 (2021).

40. Wang, Y., Bergelson, S. & Feschenko, M. Determination of Lentiviral Infectious Titer by a Novel Droplet Digital PCR Method. Hum Gene Ther Methods 29, 96–103 (2018).

41. Cooney, A.L. et al. Reciprocal mutations of lung-tropic AAV capsids lead to improved transduction properties. Front Genome Ed 5, 1271813 (2023).

